# Comparison of bacterial suppression by phage cocktails, dual-receptor generalists, and coevolutionarily trained phages

**DOI:** 10.1101/2022.07.14.500114

**Authors:** Joshua M. Borin, Justin J. Lee, Krista R. Gerbino, Justin R. Meyer

## Abstract

The evolution and spread of antibiotic resistant bacteria have renewed interest in phage therapy, the use of bacterial viruses (phages) to combat bacterial infections. The delivery of phages in cocktails where constituent phages target different modalities (e.g., receptors) may improve treatment outcomes by making it more difficult for bacteria to evolve resistance. However, the multipartite nature of cocktails may lead to unintended evolutionary and ecological outcomes. Here, we compare a 2-phage cocktail with a largely unconsidered group of phages: generalists that can infect through multiple, independent receptors. We find that both generalists and cocktails that target the same receptors suppress bacteria similarly for ~2 d. Yet a “trained” generalist phage, which previously adapted to its host via 28 d of coevolution, demonstrated superior suppression. To understand why the trained generalist was more effective, we measured the resistance of bacteria against each of our phages. We find that, when bacteria were assailed by 2 phages in the cocktail, they evolved mutations in *manXYZ,* a host inner-membrane transporter that λ uses to move its DNA across the periplasmic space and into the cell for infection. This provided crossresistance against the cocktail and untrained generalist. However, these mutations were ineffective at blocking the trained generalist because, through coevolutionary training, it evolved to bypass *manXYZ* resistance. The trained generalist’s past experiences in training make it exceedingly difficult for bacteria to evolve resistance, further demonstrating the utility of coevolutionary phage training for improving the therapeutic properties of phages.

## Introduction

The evolution of antibiotic resistant bacteria is a major threat to human health. Recent estimates suggest that >1 million deaths were attributable to antibiotic resistant infections in 2019, with mortality expected to climb to >10 million deaths in the next 30 years (Murray et al. 2022; World Health Organization, 2019). To combat the evolution and spread of resistance, many are developing evolutionarily informed strategies of antibiotic use, such as deploying drugs sequentially, in fluctuation, or at concentrations less likely to select for high levels of resistance (Imamovic & Sommer, 2013; Kim et al., 2014; Oz et al., 2014; Andersson & Hughes, 2014). In the case of life-threatening infections, clinicians often administer combinations (i.e., cocktails) of antibiotic drugs in unison, hoping to reduce bacteria’s ability to evolve resistance by forcing them into the difficult challenge of evolving resistance against multiple drugs simultaneously (Mouton, 1999; Joshi, 2011; Tamma et al., 2012). However, as bacteria have and continue to evolve resistance to the panacea of available drugs and the pipeline of drug development slows, there is growing interest in alternative ways to treat bacterial infections (Cooper & Shlaes, 2011; Tommasi et al., 2015).

Phage therapy, the use of bacterial viruses (phages) to treat bacterial infections, is a promising alternative to antibiotic drugs (Schooley et al., 2017; Chan et al., 2018; Dedrick et al., 2022). Often, phages are administered to patients in cocktails, comprised of multiple, distinct phage strains (Schooley et al., 2017; Dedrick et al., 2022). Akin to antibiotic, antiviral, and cancer combination therapies, the goal is to target multiple, distinct modalities to improve bacterial killing and prevent or delay the evolution of resistance (Gordillo Altamirano & Barr, 2021). In the context of phages, modalities constitute different infection mechanisms (e.g., the use of distinct receptors). Multiple studies of phage therapy have found that cocktails are more suppressive than individual constituent phages, especially in cases where they contain phages known to target different receptors (Tanji et al., 2004; Gu et al., 2012; Yang et al., 2020). However, increasing the complexity of multipartite phage cocktails is expected to reduce the predictability of evolutionary, ecological, and pharmacokinetic dynamics resulting from their use (Chan et al., 2013; Gordillo Altamirano & Barr, 2021). In the complex environment inside a patient, some of the phages may not be maintained at all sites of the infection, allowing bacteria to sequentially gain resistance to each constituent phage.

One unique solution to the evolution of resistance may be the deployment of “generalist” phages that infect cells through multiple, alternative receptors. Generalist phages are distinct from phages that use co-receptors to assist in adsorption, as well as from all other antimicrobials because a single genotype can kill cells via different, independent modalities, like an all-in-one cocktail. Because generalist phages are not subjected to the demographic stochasticity that affects multiphage cocktails, they may be less susceptible to resistance evolution because generalists provide a more consistent pressure on bacteria at all sites of infection.

By reciprocally adapting to changes in their hosts (coevolution), phages can evolve from single-receptor specialists into multi-receptor generalists (Meyer et al., 2012). This evolutionary capacity can be harnessed to produce generalist phages that have improved therapeutic efficacy. Previously, we demonstrated that by preemptively adapting a specialist phage to target bacterial hosts in a process called coevolutionary phage training, we could evolve generalist phages that showed 1000-fold improved bacterial suppression and delayed the evolution of resistance 14+ days compared to their phage ancestor (Borin et al., 2021). These trained generalists were much more suppressive than trained specialists, suggesting that the dominant driver of improved efficacy was the evolution to use two receptors.

These results suggest that generalist phages could be advantageous for treating bacterial infections. Yet, their use in phage therapy has not been described and no direct comparison between generalists and cocktails has been made. This is likely due to two factors: 1) practitioners often do not know the receptor(s) that their phages use (Gordillo Altamirano & Barr, 2021) and 2) generalists seem to be rare in nature. In a meta-analysis of 17 gram-positive and 64 gram-negative phages, Bertozzi Silva et al. (2016) show that the majority of phages either use a single receptor or a primary co-receptor that improves adsorption before irreversibly binding a secondary receptor. Phage T2 is an exception because it can infect using either receptor OmpF or FadL (Hantke, 1978; Black, 1988; Kortwright et al. 2020). We also determined the receptors of 17 lambdoid coliphages in our collection and found that all specialize on a single receptor (Table S1).

In this study, we evaluate the efficacy of dual-receptor phage generalists and a phage cocktail comprised of 2 specialists to suppress bacteria in vitro. To make direct, controlled comparisons, we consider highly related λ phage genotypes. These include two different generalists that both exploit host outer membrane protein receptors LamB and OmpF, as well as two specialist phages that exploit either LamB or OmpF. When we compared our cocktail with an early generalist phage (that recently evolved the ability to use 2 receptors), we find similar dynamics of bacterial suppression, suggesting treatments are equivalent. However, phage generalists that have been “trained” via coevolution with their hosts for a prolonged period were significantly more suppressive than the cocktail for 15 d. By characterizing the resistance of coevolved bacterial isolates, we find that the trained generalist is more effective because, in coevolutionary training, it evolved mutations that allow it to bypass intracellular forms of resistance.

## Materials and Methods

### Bacterial and Phage Strains

To make useful comparisons between phage cocktails and phage generalists, we leveraged a collection of highly related λ phage genotypes (Fig. 1). All phages are derived from the strictly lytic λ strain cI26 (λanc). When λanc coevolves with *Escherichia coli* B strain REL606, it repeatably evolves the ability to use a novel outer membrane protein receptor (OmpF) in addition to its native receptor LamB (Meyer et al., 2012). The early generalist phage (λegen) was isolated on day 8 of a coevolution experiment, immediately after gaining the ability to use a new receptor, OmpF, in addition to its native receptor, LamB (EvoC in Meyer et al., 2012). For our cocktail, we use one LamB specialist and one OmpF specialist (λLspec and λOspec, evolved as in Meyer et al., 2016). Because coevolutionary dynamics between λ and *E. coli* lead to dual-receptor generalists (Gupta et al., 2022a), our *specialists* were obtained by evolving λegen on hosts possessing either LamB or OmpF for 35 d. Hosts were replenished daily and therefore did not coevolve alongside the phages. A handful of mutations separate λanc, λegen, λLspec, and λOspec, with most mutations in the phages’ host recognition protein *J* (Table S2). We also compared a trained generalist phage (λtgen), isolated after 20 more days of coevolution from the same population as λegen (λtrn in Borin et al., 2021). We consider this phage “trained” because it has coevolved with its host for a prolonged period of time. During this period, λtrn evolved point mutations in genes *H, lom, Orf-401,* and *Orf-64,* as well as a recombination in *J,* and therefore differs from the other phages at more sites in the genome (Table S2).

**Figure 1.**
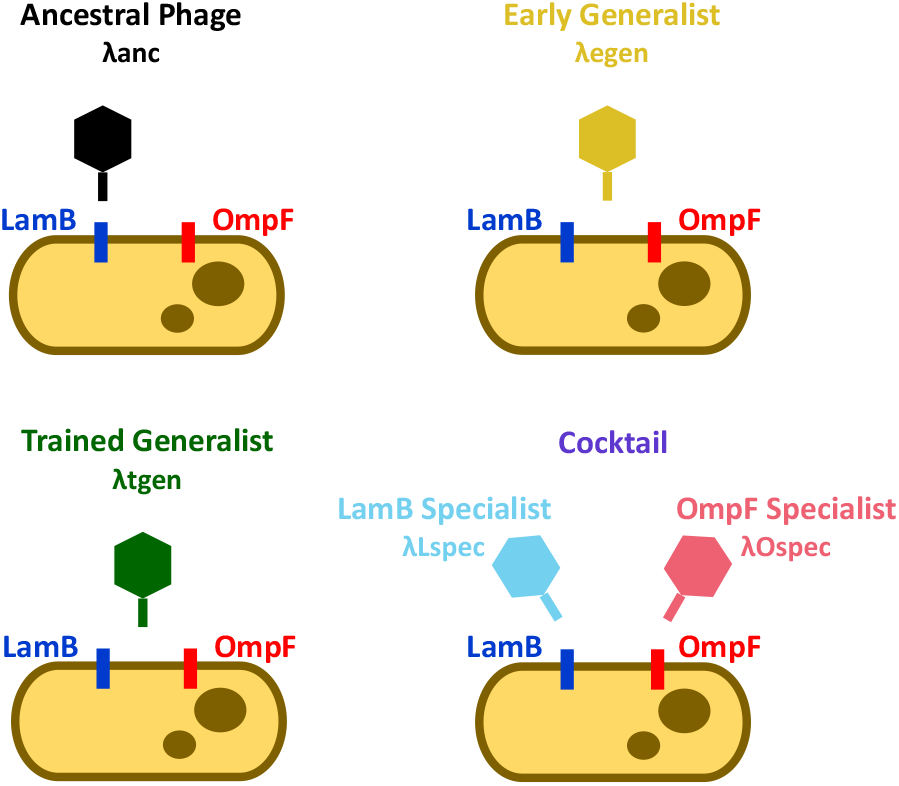
Schematic of phage strains in the study. The ancestral phage (λanc) is cI26, a strictly lytic strain of phage λ. The early generalist (λegen) is a descendant of λanc isolated after 8 d of coevolution with *E. coli.* The trained generalist (λtgen), was isolated from the same population as λegen after 20 more days of coevolution. The cocktail is comprised of a LamB specialist (λLspec) and an OmpF specialist (λOspec) which are descendants of λegen. λanc uses LamB, λegen and λtgen can use LamB and OmpF, and λLspec and λOspec use LamB or OmpF, respectively. Colors indicate phages/treatments throughout the study.

### Bacterial Suppression Experiments

To evaluate the suppressive ability of phages, we inoculated REL606 host and respective phages into 50-mL flasks with 10 mL modified M9-G (recipe in Meyer et al., 2012). Cultures were incubated at 37°C for 24 h and every day 100 μL from each population was propagated into new flasks with 10 mL of fresh media. Aliquots were removed each day to estimate bacterial and phage densities, as well as to preserve communities for later analyses. To estimate bacterial and phage densities, aliquots were serially diluted in M9-G media. Bacteria were plated on Luria-Bertani (LB) agar plates that were incubated at 37°C to count colony forming units (CFU). Phages were aliquoted in 2-μL droplets onto a soft agar overlay (LB agar except with 0.8% w/w agar) infused with ~10^8^ cells of REL606 and incubated at 37°C to enumerate plaque forming units (PFU). To preserve communities, aliquots were preserved by freezing at −70°C in 15% v/v glycerol. Experiments were initiated with the following bacterial and phage inoculums: To compare λanc, λegen, and the cocktail (comprised of λLspec and λOspec) in the first suppression experiment, ~10^7^ phages and ~10^5^ cells were inoculated into flasks. Effort was made to balance the ratio of λLspec to λOspec in the cocktail but it was inoculated at ~9:1. To compare λegen and the cocktail in the second suppression experiment, ~10^6^ phages and ~10^5^ cells were inoculated into flasks and the ratio of λLspec to λOspec in the cocktail was ~75:25. To compare λtgen and the cocktail in the third suppression experiment, ~10^6^ λtgen or ~10^7^ cocktail phages (λLspec λOspec ratio of ~75:25) were inoculated into flasks with ~10^6^ cells.

### Measuring Receptor Preference

We measured the receptor preference of phage populations by enumerating those that could use the LamB and OmpF receptors. We did this by aliquoting phages onto soft agar infused with *E. coli* K-12 Keio gene knockout collection strains (Baba et al., 2006) that were either Δ*ompF* (O^-^, strain JW0912) or Δ*lamB* (L^-^, strain JW3996). To quantify the phages’ proclivity for LamB and OmpF receptors, phage titers on LamB (O^-^ hosts) and OmpF (L^-^ hosts) were used to calculate the Specialization Index (SI), where SI = (titer_LamB_ – titer_OmpF_) / (titer_LamB_ + titer_OmpF_). SI can range from −1 to +1, indicating complete specialization on OmpF or LamB, respectively.

### Bacterial Isolation and Measuring Phage Resistance

We isolated bacteria by streaking ~2 μL of preserved, frozen communities onto LB agar plates. Plates were incubated overnight at 37°C and then colonies were isolated and streaked twice more to purify them of phage and obtain isogenic strains. Lastly, bacterial isolates were grown overnight at 37°C in LB Lennox broth and preserved by freezing. To determine when phage resistance evolved, we isolated 12 coevolved bacteria from each community across various days of the suppression experiment (~600 strains). We measured resistance by aliquoting phages onto soft agar plates infused with different isolates. By dividing the number of plaques produced on different bacterial isolates by the number of plaques formed on the ancestor (REL606), we calculated the efficiency of plaquing (EOP, a metric often used to indicate how well phages are adapted to different hosts) for each phage–host pair (~2400 pairwise EOPs).

### Survey and Test of *manXYZ* Mutations

To determine which bacterial populations and timepoints had evolved mutations in the *manXYZ* operon, we streaked one representative bacterial isolate from each population across days 1–5 on tetrazolium-mannose indicator plates (10 g tryptone, 1 g yeast extract, 5 g sodium chloride, 16 g agar, 10 g mannose per liter of water, and supplemented to a final concentration of 0.005% triphenyl tetrazolium chloride [TTC] indicator dye). Colonies formed by bacteria with mutations that disrupt mannose metabolism appear dark red and colonies without *manXYZ* mutations are pink (Burmeister et al., 2021). We then tested whether *manXYZ* mutations recapitulate the resistance we observed in coevolved bacteria from the suppression experiment by measuring the EOP of phages on Δ*manY* and Δ*manZ* Keio knockout collection strains (JW1807 and JW1808, respectively) with respect to K-12 wildtype (Baba et al., 2006).

## Results

### Suppression by the phage cocktail and early generalist

Initially, we compared the suppressive efficacy of our phage cocktail, early generalist (λegen), and their wildtype phage ancestor (λanc) (Fig. 2A). Consistent with previous studies (Borin et al., 2021), we found that the ancestral phage lost the ability to suppress bacteria in ~1 d. Both early generalist and cocktail treatments appeared more suppressive than λanc for the first 2 d. The cocktail was significantly more suppressive (p = 0.028 and p = 0.024 for days 1 and 2, respectively, Mann-Whitney U test), however we did not find statistical significance between λanc and λegen (p = 0.1 each day, Mann-Whitney U test) because sample sizes were small. In comparisons between the cocktail and early generalist, bacterial suppression was similar; both λegen and the cocktail eliminated bacteria from 1 of 3 and 2 of 6 flasks, respectively. However, the cocktail was more suppressive on day 2 (p = 0.048). Because bacterial titers were highly variable and sample sizes were small, we repeated the experiment with more replicates. (n = 10 λegen flasks and 11 cocktail flasks). On average, both λegen and the cocktail suppressed bacteria ~100-fold for the first 4 d of the experiment (Fig. 2B). There were no significant differences between treatments, suggesting that the cocktail and early generalist are equally effective at suppressing bacteria.

**Figure 2.**
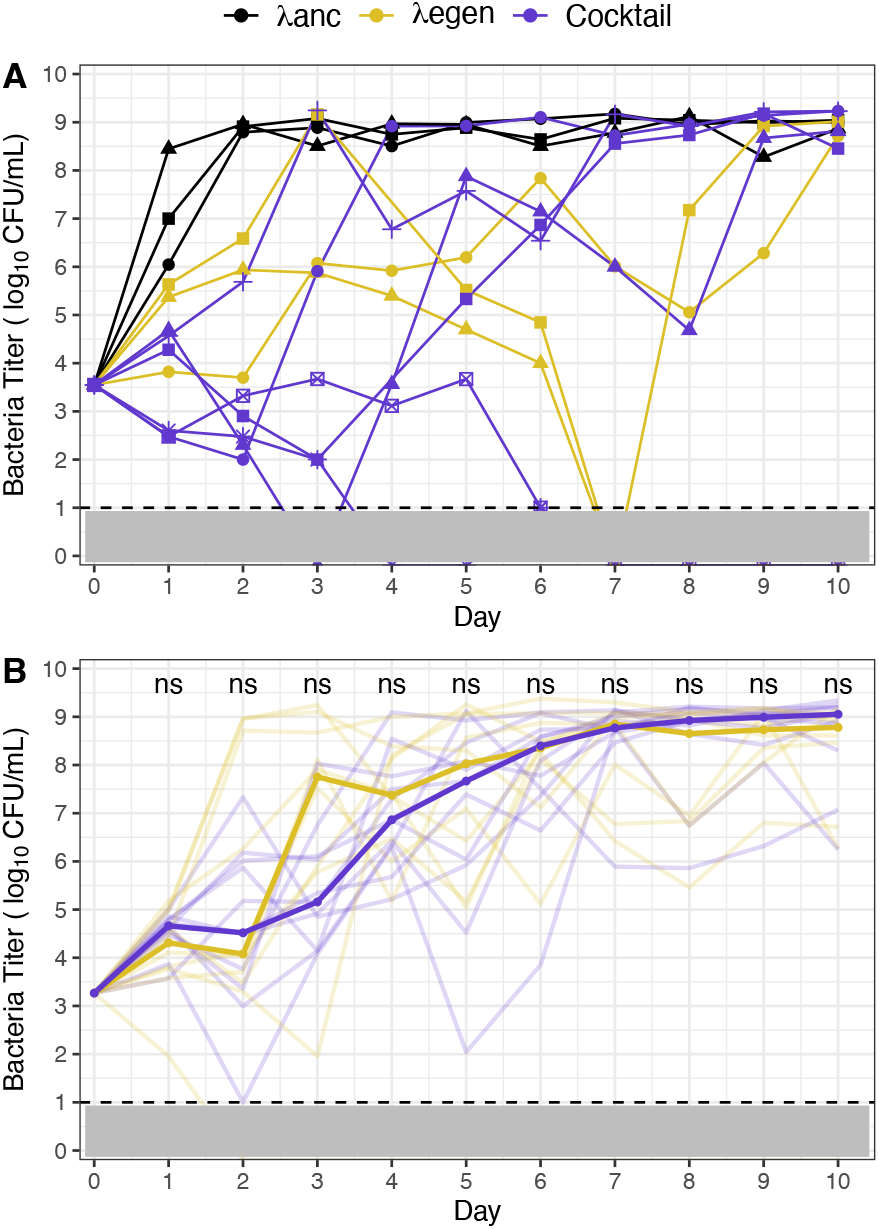
Bacterial suppression due to phage treatments. Panel A: Comparison of λanc, λegen, and cocktail treatments. Lines and shapes represent replicates (3, 3, and 6 flasks for λanc, λegen, and cocktail, respectively). Panel B: Comparison of λegen and cocktail treatments. Median bacterial titer is emboldened and trajectories of replicate flasks are translucent (10 and 11 flasks for λegen and cocktail, respectively). Mann-Whitney U tests were used to compare bacterial titer between treatments for each day. No significant differences were found (α = 0.05, ‘ns’ indicated above days). Panels A and B: λanc, λegen, and cocktail are colored black, gold, and purple, respectively. Dashed line and gray shading indicate the limit of detection (10 CFU/mL).

Combination therapies rely on the concurrent exploitation of multiple modalities to improve efficacy and delay resistance. Loss of, or asymmetries in targeted modalities can reduce the effectiveness of treatment. Because the cocktail is a multipartite treatment comprised of 2 distinct specialist phages, we investigated how its composition (and therefore preference for different receptors) differed from the early generalist treatment, which is comprised of a single genotype. In addition to estimating the overall phage titer each day, we measured the number of phages that could infect using LamB or OmpF receptors by aliquoting the community phage lysate on hosts lacking either OmpF (O^-^) or LamB (L^-^), respectively. We then calculated the Specialization Index (SI), which quantifies phages’ preference on a scale from 1 (LamB specialization) to –1 (OmpF specialization). We hypothesized that over time the SI of the cocktail would either H1) fix at –1 or 1, coinciding with fixation of a specialist genotype, H2) occupy intermediate values suggesting a balance between the specialist phages or H3) fluctuate between –1 and 1 in a negative frequency-dependent manner. Support for H1 or H3 associated with a loss of bacterial suppression could suggest that efficacy is lost due to a loss of, or asymmetry in receptor use.

We find that SI values of the cocktail fluctuated between 1 and –1, indicating that the composition of the cocktail alternated between λLspec and λOspec phages (Fig. 3A). These oscillations, which initially favor LamB specialization (SI ~ 1), give way to OmpF specialization (SI ~ −1) as the cocktail maintains suppression, and then return to SI ~ 1. In contrast, the SI of the early generalist maintained intermediate values (Fig. 3B). When we repeated the experiment with more replicate populations (Fig. 2B), we found similar dynamics, albeit with more variable periods of oscillation (Fig. S1). We did not observe any relationship between the predominance of λLspec or λOspec and the cocktail’s ability to maintain suppression. For example, in cocktail population 5, the rapid shift to LamB specialization correlates with lower suppression, whereas in population 4, the shift to LamB specialization coincides with increased suppression (Fig. S1). Altogether, these results show that fluctuations in the composition and receptor preference of the cocktail helped maintain phages throughout the experiment and possibly prolonged the cocktail’s effectiveness.

**Figure 3.**
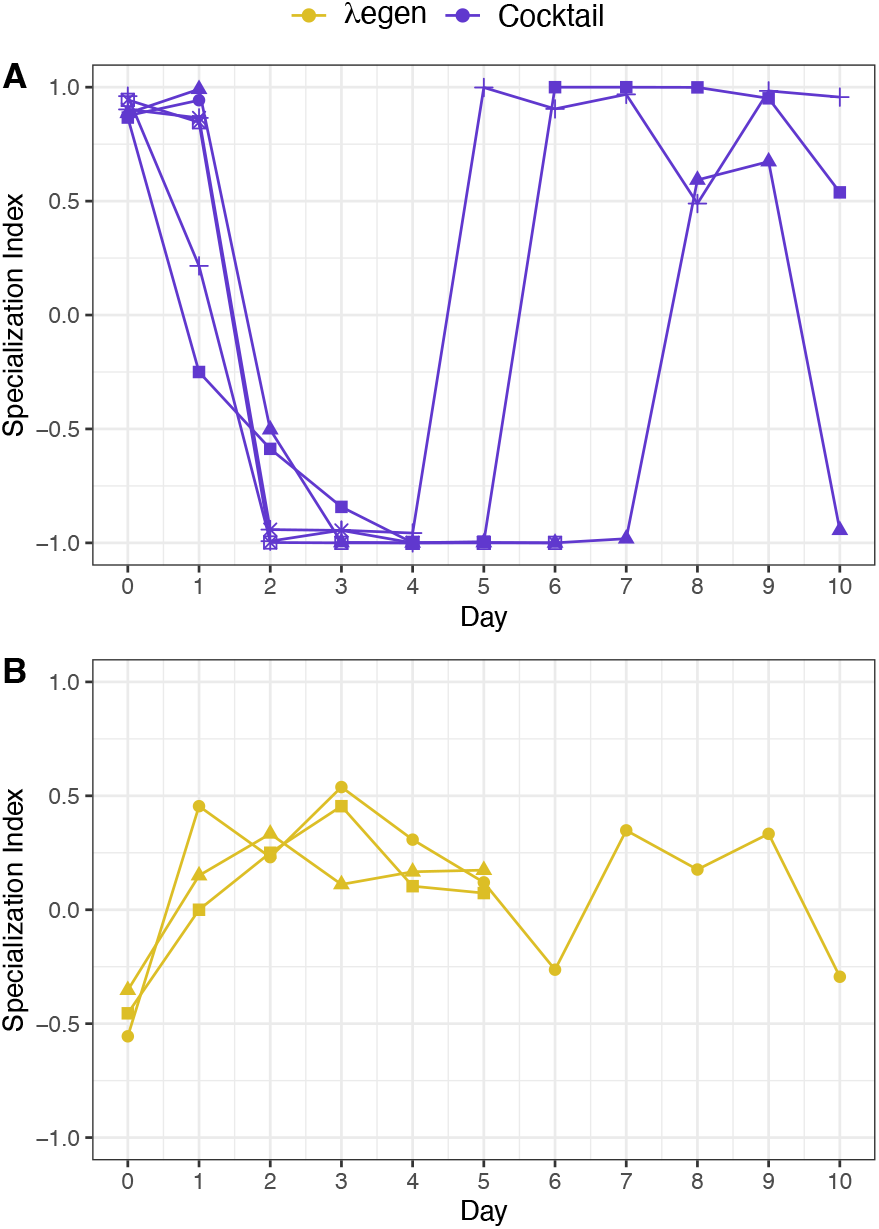
Preference of λegen and cocktail treatment for LamB and OmpF receptors over days of the first suppression experiment (Lines and shapes represent replicates and correspond with those in Fig. 2A). The cocktail (A) is colored purple and λegen (B) is gold. Receptor preference was quantified using a Specialization Index (SI) where SI = (titer_LamB_ – titer_OmpF_) / (titer_LamB_ + titer_OmpF_). SI ranges from 1 (LamB specialization) to –1 (OmpF specialization).

### Suppression by the phage cocktail and trained generalist

In previous work, we demonstrated that the ability to infect through 2 distinct receptors substantially improves bacterial suppression (Borin et al. 2021). We did this by comparing phages that were “trained” via coevolution with their host for 20 d. Half of the phages we compared had evolved the ability to use an additional receptor during training and were far more suppressive (~1000-fold) than the phage ancestor. However, the other half of the phages in our comparison did not evolve to use a new receptor and showed only meager improvements in suppression. This led us to conclude that the ability to infect through 2 receptors (which evolved as a result of phage training) drastically improves suppressive efficacy. However, when we compared λegen and the cocktail (above), which both exploit 2 receptors, neither seemed as suppressive as the trained dual-receptor phages from the previous study.

To investigate this discrepancy, we compared a trained generalist phage (λtgen) against our cocktail in another suppression experiment. λtgen was significantly more suppressive for 15 d, and bacteria were driven extinct in 3 of 5 λtgen flasks (Fig. 4), clearly showing that the trained generalist was more suppressive than the early generalist and phage cocktail. Again, we found that the cocktail fluctuated between LamB and OmpF specialization, whereas the trained generalist maintained intermediate SI values like the early generalist. (Fig. S2). Because λegen, λtgen, and the cocktail all use the same 2 receptors, these results suggested that λtgen has some other property that makes it more effective at suppressing bacteria.

**Figure 4.**
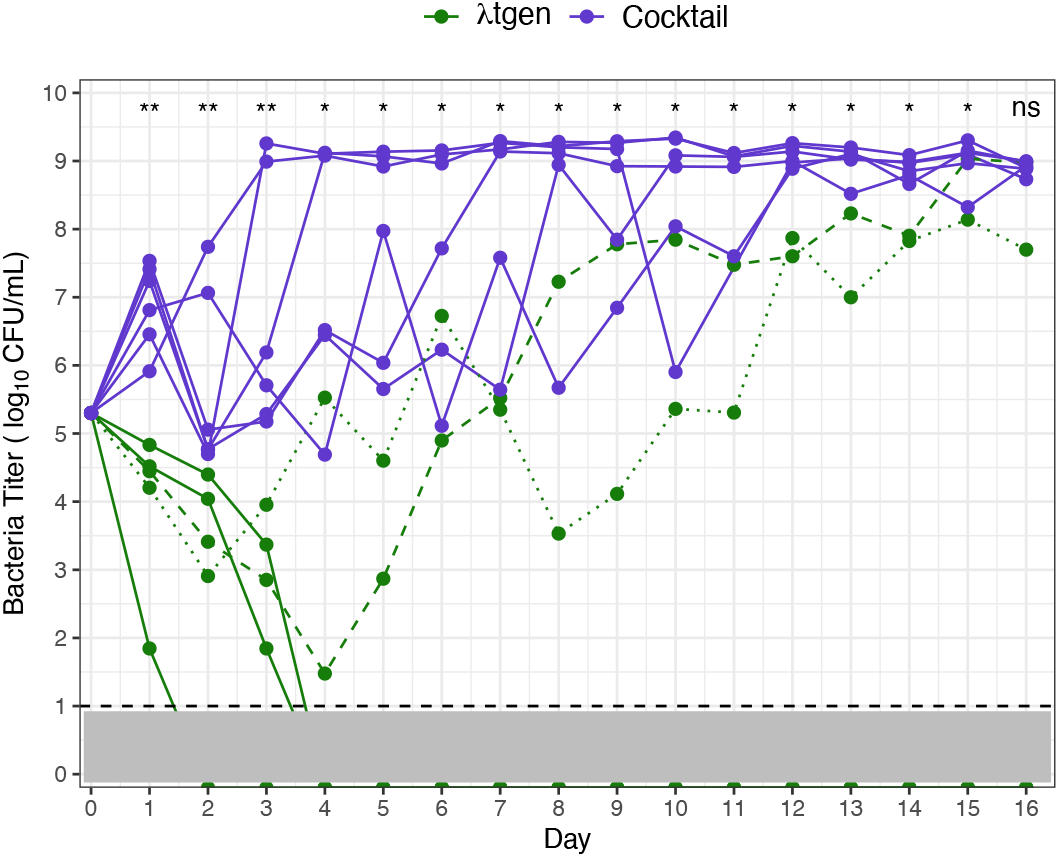
Bacterial suppression in λtgen (green) and cocktail (purple) treatments. Lines represent replicate flasks (n = 5 with λtgen and 6 with the cocktail). Bacteria were driven extinct in 3 λtgen flasks; For λtgen flasks where bacteria survived, dashed and dotted lines indicate populations 1 and 2, respectively. The dashed line and gray shading denote limit of detection. Significant differences are indicated above days, calculated via Mann-Whitney U test (not significant [ns] p > 0.05, * p < 0.05, ** p < 0.005).

### Evolution of resistance

To investigate why the trained generalist was able to suppress bacteria for so much longer than the cocktail, we first measured *when* bacteria evolved resistance in each treatment. We isolated 12 coevolved bacteria from various timepoints of each population in the suppression experiment. Then, we measured how resistant each coevolved bacterial isolate was to each of our phages using efficiency of plaquing (EOP, a metric of how well phages’ infecs different hosts). By aliquoting all of our phages (λtgen, λLspec, λOspec, λegen) on each host, we measured how resistant bacteria were to phages within their treatment, as well as to phages outside of their treatment. For example, we measured the resistance of bacteria in the λtgen treatment to λtgen (within treatment), as well as to λegen, λLspec, and λOspec (outside of treatment).

Consistent with previous work, we found that λtgen maintained suppression because it delayed the evolution of phage resistance (Fig. 5A). Of the 2 surviving λtgen bacterial populations, high levels of resistance did not evolve for >10 days and, in population 2, high levels of resistance to λtgen never evolved. Because the pathway by which REL606 evolves resistance to λtgen has previously been characterized (Borin et al., 2021), we focused our investigation on how bacteria evolved resistance in the cocktail treatment.

**Figure 5.**
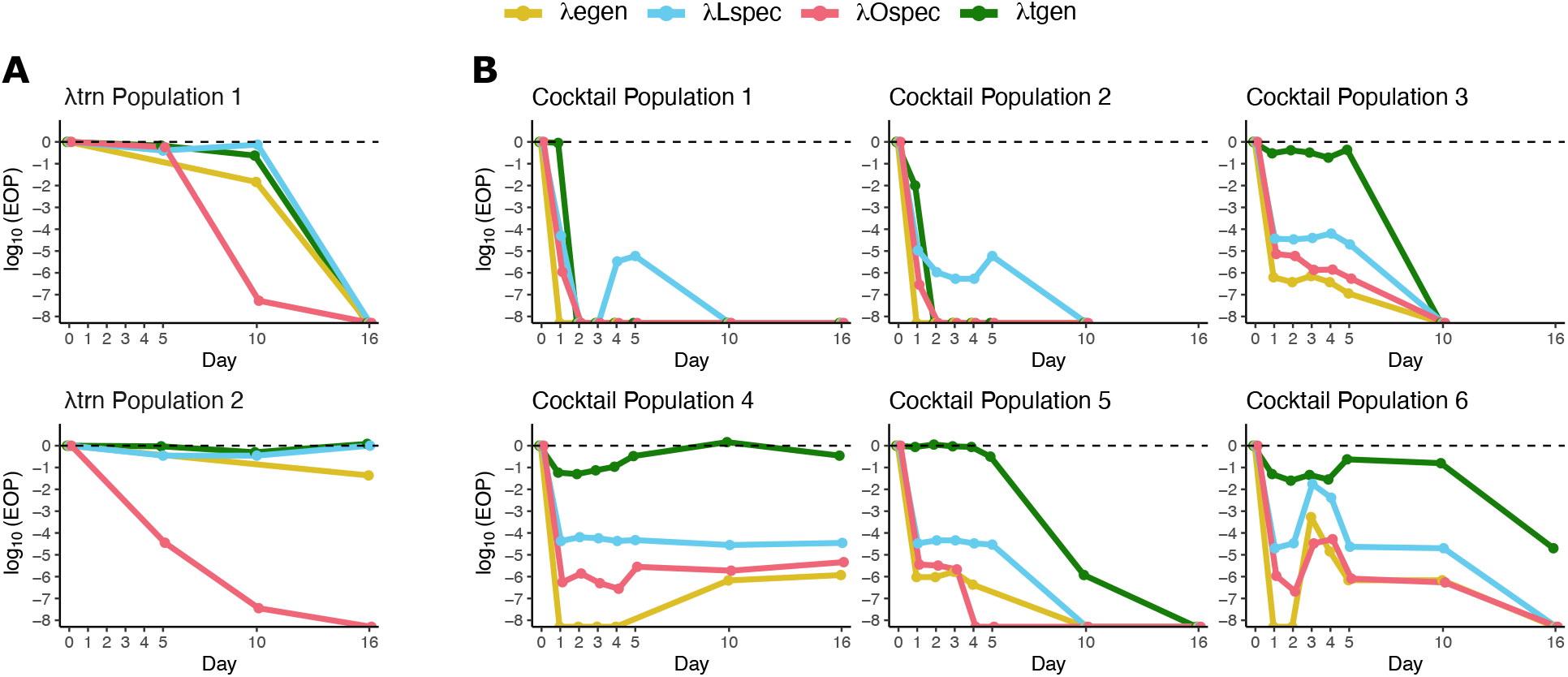
Median resistance of n=12 coevolved bacterial isolates across various days of the suppression experiment against our panel of 4 phages. Resistance of bacteria to λegen, λLspec, λOspec, and λtgen is indicated by gold, blue, red, and green lines, respectively. Resistance was measured by calculating the efficiency of plaquing (EOP = plaques on coevolved bacteria / plaques on bacterial ancestor). EOP is presented on a log_10_ scale where the dashed line indicates EOP = 1 (where bacterial isolates are as sensitive as the ancestor). Points along the x-axis indicate where phages were unable to form plaques on bacteria from that timepoint (EOP = 0).

Bacteria in all cocktail populations evolved high levels of resistance (EOP <10^-4^) to both λLspec and λOspec within 1 day (Fig. 5B), explaining why the cocktail treatment lost efficacy much earlier than λtgen. We also noticed that in all populations, resistance to λLspec and λOspec coincided with high levels of resistance to λegen. Yet, the bacteria in 4 of 6 cocktail populations did not evolve concomitant levels of λtgen resistance, indicating that resistance mutations that evolved early against the cocktail also conferred high levels of resistance to λegen, but not to λtgen. These results also suggest that λtgen may be more suppressive because it is less susceptible to easily acquired mutations that confer cocktail (λLspec, λOspec) and λegen resistance.

Next, we investigated *how* bacteria evolved resistance. Because all of our treatments, including λtgen, exploit the same 2 receptors, it seemed unlikely that cocktail resistance would be explained by mutations in LamB and OmpF. Therefore, we considered other mechanisms that might provide resistance against the cocktail and early generalist phages but not to λtgen. A large body of work has shown that the *E. coli* mannose transporter, encoded by the *manXYZ* operon (formerly *ptsM,* a phosphoenolpyruvate-dependent phosphotransferase system), which is embedded in the inner, cytoplasmic membrane, is used by phage λ to move its DNA across the periplasmic space and into the cell for infection (Scandella & Arber, 1976; Casjens & Hendrix, 2015). Additionally, the evolution of *manYZ* mutations have been found in previous coevolution experiments between λ and *E. coli* (Meyer et al., 2012; Burmeister et al., 2021; Gupta et al. 2022b). Therefore, we used tetrazolium-mannose indicator plates to survey bacterial isolates from days 1–5 of all λtgen and cocktail populations for *manXYZ* mutations. We found *manXYZ* mutants in every isolate from the cocktail treatment and none in λtgen populations. The ubiquity of *manXYZ* mutations in cocktail populations by day 1 suggests that it is the primary cause of phage resistance and loss of bacterial suppression. Moreover, the lack of *manXYZ* mutants in λtgen populations suggest that they did not arise because these mutations may not be effective for λtgen-resistance.

To directly test whether mutations in *manXYZ* provide protection from infection, we measured the resistance of Δ*manY* and Δ*manZ* strains (that do not have any other resistance mutations) against each of our phages. We find that the resistance profiles of Δ*manY* and Δ*manZ* strains closely resemble the resistance of coevolved bacteria from the cocktail treatment (Fig. 6); Δ*manY* and Δ*manZ* hosts are highly resistant to λLspec, λOspec, and λegen (EOP = 0.05 – 3 × 10^-5^) and less resistant to λtgen (EOP ~ 0.25). The relative resistance of Δ*manY* and Δ*manZ* strains also matches patterns from the cocktail treatment; they are most resistant (lowest EOP) to λegen, followed by λOspec, λLspec, and then λtgen. Altogether, these results further support that, when treated with the cocktail, bacteria rapidly evolved resistance via mutations in *manXYZ* and also that these mutations do not grant high levels of resistance to λtgen.

**Figure 6.**
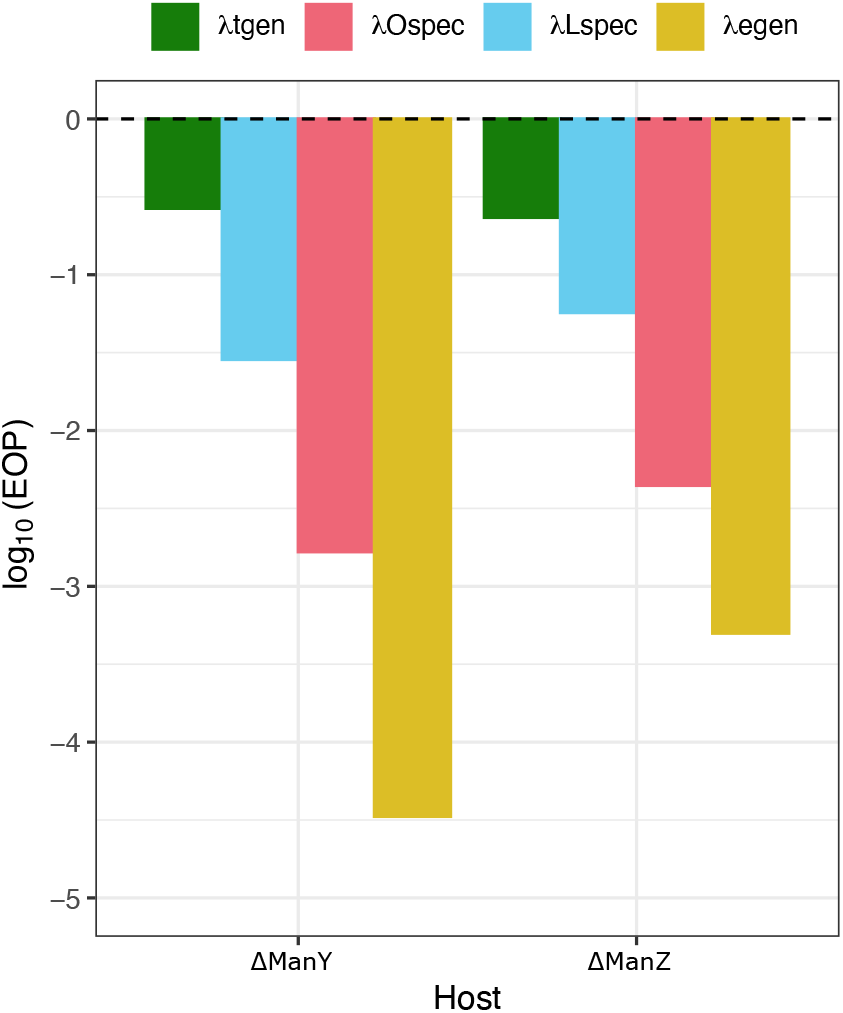
Resistance of *E. coli* K-12 ΔManY and ΔManZ knockout bacteria to our panel of 4 phages. Resistance to λegen, λLspec, λOspec, and λtgen is indicated in gold, blue, red, and green, respectively. Resistance was measured by calculating the efficiency of plaquing (EOP = plaques on knockout bacteria / plaques on bacterial ancestor). EOP is presented on a log_10_ scale where the dashed line indicates EOP = 1 (focal bacteria is as sensitive as the ancestor).

Previous studies of phage λ offer insight into how λtgen is able to bypass *manXYZ* resistance mutations. Researchers have found that adaptive mutations in λ genes *V* (major tail subunit) or *H* (tape measure protein and tail component) can allow it to infect resistant *manXYZ* mutants (Scandella & Arber, 1976; Esquinas-Rychen & Erni, 2001). In the case of our phages, λOspec and λtgen each have a unique *H*_mutation (Table S2). λOspec’s *H* mutation clearly does not allow it to overcome *manXYZ* resistance. However, for λtgen, the ability to infect *manXYZ* mutants that are resistant to λegen and the cocktail may be due to its *H* mutation, due to a different mutation (λtgen also has mutations in *lom, Orf-401,* and *Orf-64,* for which functions are not fully understood), or due to a combination of mutations. Yet these results explain why *manXYZ* mutants did not evolve in the λtgen treatment; *manXYZ* mutations do not confer high levels of resistance to λtgen because λtgen has already evolved mutations for counter-resistance. These counter-measures allowed λtgen to remain effective while the phage cocktail lost suppression because of easily acquired *manXYZ* resistance mutations.

## Discussion

In phage therapy, multi-phage cocktails are often used to improve treatment outcomes. Ideally, cocktails are comprised of phages that work additively or synergistically to kill target bacterial pathogens. For example, phages that target different host receptors can make it more difficult for bacteria to evolve resistance (Tanji et al., 2004; Gu et al., 2012; Yang et al., 2020). Previous work has also demonstrated that generalist phages, which infect using multiple, distinct receptors, can also improve bacterial killing and stem the evolution of resistance (Borin et al., 2021). Here, we leveraged a collection of highly related λ phage strains to make novel comparisons between cocktails and generalists.

We conducted initial comparisons between a cocktail, comprised of two specialist phages (λLspec and λOspec) and an early generalist phage (λegen) that recently evolved the ability to use two receptors. We found that the cocktail and λegen treatments were both more suppressive than their phage ancestor, supporting previous studies demonstrating that phage therapeutics that target multiple receptors improve efficacy (Tanji et al., 2004; Gu et al., 2012; Yang et al., 2020). We also found that the cocktail and λegen showed similar levels of suppression. The composition of the cocktail fluctuated throughout the experiment. However, we found no link between fluctuations and a loss or gain in suppressive efficacy. Our results suggest that similar outcomes in therapy may be achieved, regardless of whether the targeted host receptors are exploited by a single generalist or multiple specialists. However, caution should be used extrapolating these results to animal host infections. On the one hand, kill-the-winner dynamics that promote negative frequency-dependence and maintain constituent phages at the site of infection could improve therapeutic outcomes (Maslov & Sneppen, 2017). However, the fact that fluctuations arose in a simplified flask experiment may be concerning for in vivo applications if spatial heterogeneity and environmental complexity exacerbate asymmetries in the treatment. Future studies should evaluate whether the inherent compositional differences between phage cocktails and phage generalists lead to diverging outcomes when administered for in vivo phage therapy.

We also compared our cocktail against a trained generalist (λtgen) that previously demonstrated strong suppressive capabilities (Borin et al., 2021). Although both of our treatments use the same two receptors, λtgen was significantly more suppressive. It took bacteria >10 days to evolve high levels of resistance to λtgen, whereas bacteria evolved resistance to the cocktail in 1 day. By measuring how bacteria evolved resistance in each treatment, we discovered why the cocktail and λegen performed similarly, as well as why the trained generalist was superior.

Bacteria evolved resistance to both phages in the cocktail by acquiring mutations in *manXYZ*, a mannose transporter that λ uses to traffic its DNA across the periplasm and into the host cell (Esquinas-Rychen & Erni, 2001). These *manXYZ* mutants were also resistant to the early generalist, explaining why the efficacy of our cocktail and λegen treatments were similar. When besieged by phages targeting 2 different receptors, bacteria evolved cross resistance by altering an intracellular pathway instead of mutating each receptor separately. This blocked infection, regardless of the receptor(s) used by the phage. In this study we used highly related λ strains to control for factors apart from receptor use. This could have made it easier for bacteria to evolve cross resistance. Cocktails comprised of more distantly related phages may reduce the likelihood of cross resistance, however these results highlight the importance of determining whether cross resistance mutations are easily acquired, even when phages target different receptors (Abedon et al., 2021).

The trained generalist was superior to all other treatments because *manXYZ* mutations were not effective to block λtgen infection. Previously, we concluded that coevolutionary training improved phages by selecting for those that had evolved to use 2 receptors (Borin et al., 2021). It is now clear that training also selects for phages that experience and evolve to bypass other forms of resistance, like *manXYZ*, thereby producing phages that were far less susceptible to the evolution of resistance. Because mutations to counter *manXYZ* resistance evolved after the appearance of dual-receptor generalists (λegen lacks *manXYZ* counter resistance), our results suggest that it is advantageous to continue training phages after they have acquired novel functions, in order for them to experience and counter new forms of resistance in their hosts.

Altogether our results demonstrate strong support for the use of multi-receptor targeting phage cocktails and generalists for therapy. However, we believe that practitioners should determine whether target bacteria can readily evolve cross resistance, as this may rapidly cause treatments to fail. We also find more support for coevolutionary phage training, as it produces phages that bypass mechanisms of cross resistance and drastically improves efficacy. We find that coevolutionary training is a particularly powerful approach because it employs the natural algorithm of evolution. Through coevolution with their hosts, phages like λtgen “solve” evolutionary challenges that were neither known nor anticipated by their trainers.

## Acknowledgments

We thank the labs of Ry Young, Sherwood Casjens, and Carolin Wendling for sending us lambdoid phages. This study benefitted from useful discussion with all members of Justin R. Meyer and Sergey Kryazhimskiy’s laboratories. The work herein was supported by the National Science Foundation (award 1934515) and the National Institutes of Health Cell and Molecular Genetic Training Program (T32GM007240).

