## Supplemental Materials for "Comparison of bacterial suppression by phage cocktails, dual-receptor generalists, and coevolutionarily trained phages"

Table S1. Host receptors used by 17 lambdoid phage strains. Phage lysates were aliquoted onto soft agar plates infused with *E. coli* K-12 wildtype or wildtype-derived knockout strains from the KEIO gene knockout collection. We determined the phages' receptor by whichever knockout host it failed to infect (form a zone of lysis on). For example, phage λ is unable to lyse K-12 *ΔlamB*, indicating that LamB is its receptor.

| **Phage Strain** | **Host Receptor** |
|---|---|
| Lambda (λ) | LamB |
| Φ21 | LamB |
| HK97 | LamB |
| HK629 | LamB |
| HK630 | LamB |
| Φ434 | OmpC |
| mEpX1 | FhuA |
| mEpX2 | FhuA |
| HK022 | FhuA |
| HK140 | FhuA |
| Φ80 | FhuA |
| mEp043c | FhuA |
| mEp213 | FhuA |
| mEp234 | FhuA |
| mEp235 | FhuA |
| mEp237 | FhuA |
| mEp390 | FhuA |

Table S2. Genomic differences between our λ ancestor (λanc) and the other phages in the study. "X" indicates that the mutation is present. For genomic differences between λanc and the λ reference (GenBank: NC_001416) see Meyer et al. 2012. All mutations are nonsynonymous except the recombination in λtgen which contains synonymous and nonsynonymous mutations. The recombination occurred between λ and a relict prophage in the genome of REL606 during a coevolution experiment (Meyer et al., 2012; Borin et al., 2021).

| **Position** | **Mutation** | **Gene** | **λegen** | **λLspec** | **λOspec** | **λtgen** |
|---|---|---|---|---|---|---|
| 11,451 | C → T | *H* | | | | X |
| 11,828 | A → G | *H* | | | X | |
| 17,049 – 18,297 | Recombination | *J* | | | | X |
| 18,492 | C → A | *J* | X | X | X | |
| 18,503 | C → T | *J* | | | X | X |
| 18,537 | C → A | *J* | | | X | |
| 18,538 | A → G | *J* | X | X | X | X |
| 18,589 | C → A | *J* | | X | | |
| 18,731 | C → A | *J* | | | X | |
| 18,734 | T → C | *J* | X | X | | |
| 18,814 | C → T | *J* | | | X | X |
| 18,823 | G → A | *J* | X | X | X | X |
| 18,825 | T → A | *J* | X | X | X | X |
| 18,868 | A → T | *J* | | | | X |
| 19,260 | T → C | *lom* | | | | X |
| 20,661 | A → G | *Orf-401* | | | | X |
| 39,394 | A → G | *S/S'* | | X | | |
| 45,176 | $(G)_{5\rightarrow6}$ | *Orf-64* | | | | X |

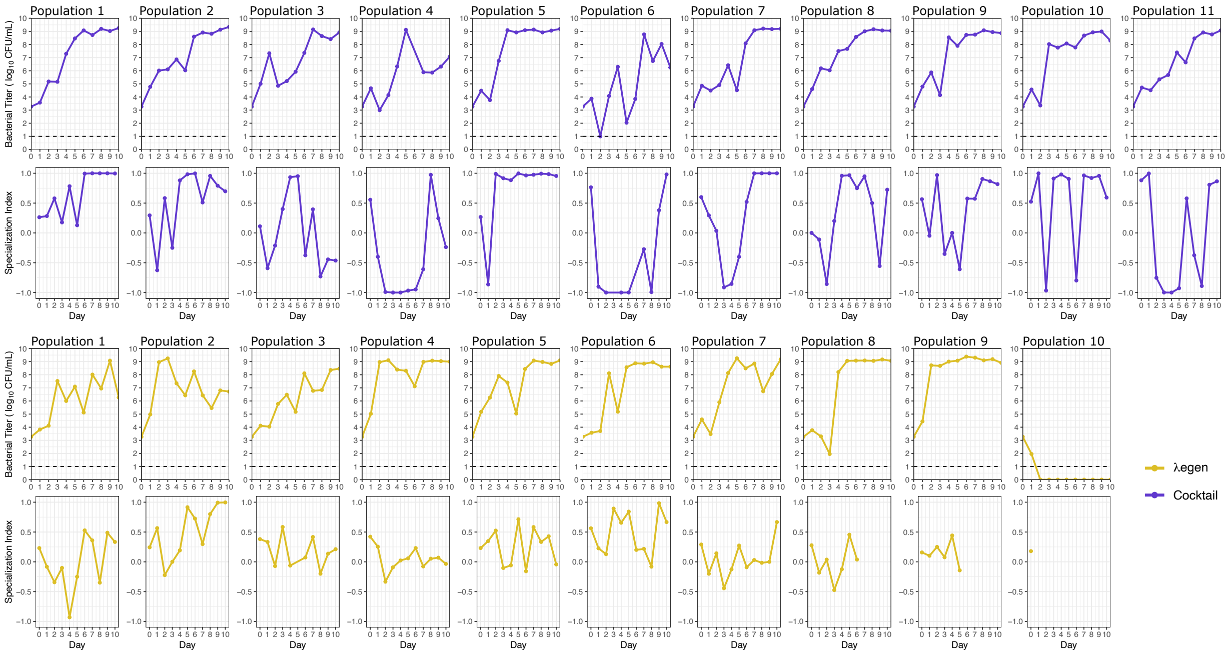


Figure S1. Bacterial titer and phage Specialization Index (SI) throughout the highly replicated suppression experiment. Lines are colored with respect to phage treatment; cocktail is purple and λegen is gold. Within each treatment, the top row shows bacterial titers and the bottom row shows SI over the 10-d experiment. Each replicate population is plotted separately, from left to right (n = 11 and 10 replicates for the cocktail and λegen treatments, respectively). In bacterial titer panels, the dashed line indicates the limit of detection (10 CFU/mL).

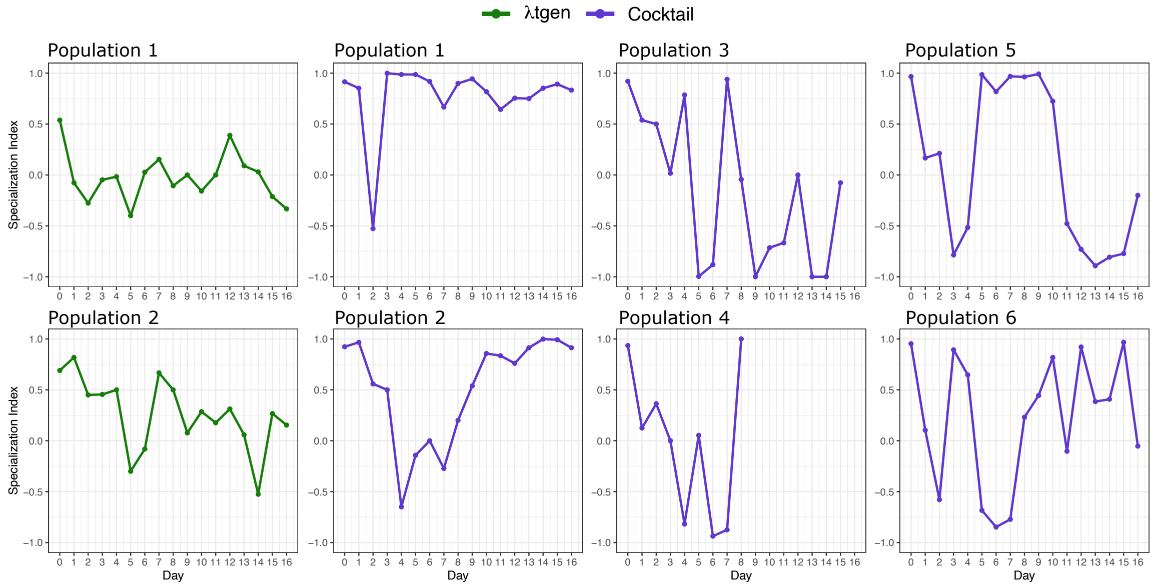


Figure S2. Phage Specialization Index (SI) from the suppression experiment comparing λtgen and the cocktail. Lines are colored with respect to phage treatment (λtgen is green and the cocktail is purple) and each replicate population is plotted separately.
